# Larger Pre-Speech Auditory Modulation Is Associated With Smaller Corrective Responses To Auditory Errors

**DOI:** 10.64898/2026.10.01.756086

**Authors:** Alexis Basciano, Sara-Ching Chao, Corianne Rogalsky, Ayoub Daliri

## Abstract

**Purpose:** During speech planning, the brain predicts the auditory consequences of its movements and modulates the auditory system—a process called pre-speech auditory modulation (PSAM). We investigated the link between PSAM and behavioral error correction and determined how aging influences these mechanisms.

**Methods:** Seventeen young adults (18–32 years) and 13 older adults (50–85 years) performed a speaking task involving auditory feedback perturbations and a silent reading control. We measured electrophysiological responses (N1/P2) to auditory probes during speech planning and correlated these modulations with the magnitude of participants’ subsequent behavioral corrective responses.

**Results:** Both groups showed significant modulation of N1 and P2. However, N1 modulation correlated with corrective responses only in young adults. Older adults showed reduced behavioral corrections and smaller, delayed P2 components. Despite these age-related behavioral differences and the loss of the modulation-correction correlation, older adults maintained modulation magnitudes comparable to those of younger adults.

**Conclusions:** N1 modulation serves as a neural index of the brain’s predictive processes during speech planning. Aging appears to selectively disrupt the integration of these signals into motor output rather than the internal generation of the predictions themselves.

---

Contemporary theories of speech motor control posit that the brain integrates feedforward and feedback control mechanisms to produce accurate and intelligible speech (Daliri, 2021; Guenther, 2016; Houde & Nagarajan, 2011; Parrell & Houde, 2019). The feedforward mechanism generates motor commands to achieve the desired auditory goals. As the brain prepares motor commands, it predicts the auditory consequences of those commands. By relying on predictive processes during planning, the feedforward mechanism optimizes motor commands to ensure stable, low-variability output (e.g., Daliri et al., 2025). During production, the feedback mechanism monitors the auditory feedback to ensure the accuracy of speech output. The feedback system relies on predictive processes to detect errors in speech output (i.e., the difference between prediction and auditory feedback) and generate “corrective” responses (Daliri et al., 2020; Parrell et al., 2017; Sreedhar & Daliri, 2026; Tourville et al., 2008). The brain also evaluates errors to determine their relevance and source. Then, the brain uses its evaluation to “adapt” its feedforward motor commands to achieve its desired goals in subsequent movements more accurately (Daliri & Dittman, 2019; Houde & Jordan, 1998; Lametti et al., 2014; Villacorta et al., 2007). A higher reliance on feedback may lead to more error sensitivity and larger motor responses to errors (i.e., larger corrective and adaptive responses) (e.g., Shadmehr & Mussa-Ivaldi, 2012). Overall, the brain relies on prediction and auditory feedback to (1) optimize and adapt its feedforward motor commands and (2) generate feedback-driven corrective responses. The relative weighting of prediction versus feedback determines a speaker’s motor stability and their sensitivity to sensory errors.

Predictive processes are frequently examined through sensory modulation, a phenomenon in which brain responses to self-generated sensory stimuli are significantly modulated (suppressed) relative to identical stimuli from an external source (Hughes et al., 2013; Waszak et al., 2012). This modulation occurs because the brain predicts the sensory consequences of its own movements, “canceling out” expected inputs to prioritize novel external information (Bays et al., 2005; Blakemore et al., 1998). While traditionally studied during movement, research now shows that these predictive processes are also active during motor planning (Eliades & Wang, 2008; Schneider et al., 2014; Seki & Fetz, 2012). For instance, by comparing neural (and perceptual) responses to auditory probes during speech planning versus silent reading, we demonstrated in a series of studies that auditory (and somatosensory) responses are modulated (suppressed) even before speech onset (Daliri & Max, 2015b, 2015a, 2016, 2018; McGuffin et al., 2020; Merrikhi et al., 2018). We recently proposed that this pre-speech auditory modulation (PSAM) serves as a neural index for the predictive processes involved in feedforward motor preparation (Li et al., 2024, p. 202). Because the brain relies on these predictions to optimize motor commands, a more robust predictive system should result in greater trial-to-trial stability. Supporting this suggestion, our analysis of the early, feedforward-dominant window of speech revealed that lower modulation levels (i.e., less suppression) correlated with increased trial-to-trial formant variability. This relationship is further evidenced in clinical populations: individuals who stutter—a disorder linked to inefficient predictive processes (Chang & Guenther, 2020; Daliri et al., 2014, 2025; Max & Daliri, 2019; Neef & Chang, 2024)—exhibit both reduced PSAM and heightened speech variability (Daliri et al., 2025; Daliri & Max, 2015b; Jackson et al., 2016; Smith & Kleinow, 2000). Collectively, these findings establish PSAM as a robust neural correlate of the predictive mechanisms required to optimize feedforward mechanisms.

Beyond motor planning, predictive processes are essential for feedback control mechanisms active during speech production (Behroozmand et al., 2025; Daliri et al., 2020; Parrell et al., 2017; Sreedhar & Daliri, 2026; Tourville et al., 2008). The brain uses these predictive processes as a template to detect discrepancies, allowing it to generate rapid corrective responses—typically within 100–200 ms of an auditory error. These responses serve as a primary behavioral measure of the brain’s online error-correction capabilities. To investigate these processes, researchers often employ “compensation paradigms,” which introduce near-real-time auditory feedback perturbations during speech. Our recent work (Daliri et al., 2020; Merrikhi et al., 2025; Sreedhar & Daliri, 2026) using this paradigm demonstrated that the feedback control mechanism is highly context-sensitive: corrective responses scale inversely with perturbation magnitude and are significantly more robust when errors cross phonetic categories (e.g., shifting /ɛ/ toward /æ/). The link between predictive processes and the feedback control mechanism is further evidenced in individuals who stutter. This population, characterized by inefficiencies in predictive processes, exhibits markedly reduced corrective responses to auditory perturbations (Cai et al., 2012; Frankford et al., 2022; Loucks et al., 2012). However, while both neural and behavioral markers of these systems have been studied independently, the direct relationship between PSAM—a neural index of prediction—and corrective responses—the behavioral measure of the feedback control mechanisms—remains to be elucidated.

The primary objective of the current study was to elucidate the relationship between PSAM and corrective responses to auditory errors. Furthermore, as the neural mechanisms supporting predictive processes undergo significant age-related changes (Bornkessel-Schlesewsky et al., 2015; Chan et al., 2021; Moran et al., 2014), a secondary objective of this study was to characterize PSAM and corrective responses in both young and old adults. To address these goals, we measured PSAM by comparing neural activity during speaking and silent reading conditions. We integrated these measures into a single task by modifying the speaking condition to include real-time auditory perturbations that shifted the formants of /ɛ/ toward /æ/. We hypothesized that if feedback control mechanisms rely on internal predictive processes, then the magnitude of PSAM would correlate with the corrective responses. Furthermore, we predicted that if aging impacts the neural substrates of prediction, then the relationship between PSAM and corrective responses would differ between young and old adults.

## Method

### Participants

We recruited two groups of participants: 17 young adults (4 males; Age 23.7 ± 4.1 years, 18–32 years) and 13 older adults (7 males; Age 68.4 ± 9.1 years, 50–85 years). All participants had normal or corrected-to-normal vision. All participants were native speakers of American English and had no current or past neurological, psychological, or communication disorders. Additionally, all participants had normal hearing thresholds (less than 25 HL) at all octave frequencies from 250 Hz to 4000 Hz in both ears (ASHA, 1997). All older adults completed the Mini-Mental Status Examination (MMSE; (Folstein et al., 1975)) to detect and exclude individuals with any preexisting cognitive impairment (all MMSE scores were 24 or higher). Arizona State University’s Institutional Review Board reviewed and approved all aspects of the experimental and recruitment procedures for this study.

### Apparatus

For the duration of the experimental session, participants were seated ∼4 feet away from a 24-inch monitor (Alienware, 240 Hz refresh rate), inside a sound-attenuated booth (Acoustic Systems, Inc.). A mounted microphone (SM58, SURE) was positioned 15 cm away from the participant’s mouth, at a 45-degree angle relative to the participant. Throughout the experimental session, a control computer presented instructions and target words on the display monitor. Once participants read the words aloud, the speech signal was collected via the microphone, amplified (TubeOpto 8), and sent to the control computer by an audio interface (MOTO 8pre USB). The audio interface output was amplified (S-phone) and binaurally delivered to the participant via insert earphones (ER2, Etymotic Research).

For presenting visual and auditory stimuli, we used the Psychtoolbox software (Kleiner et al., 2007) running within the MATLAB environment in Windows 11. Visual stimuli were rendered on the monitor at 240 Hz with a resolution of 1920 × 1080 pixels. Auditory stimuli were presented using the audio interface’s native driver (MOTU ASIO) at a 48000 Hz sampling rate. We also used Audapter for near-real-time formant tracking and shifting (Cai, 2015). We used the following Audpater parameters that resulted in an input-to-output delay of ∼18 ms: LPC order of 17 for males and 15 for females; sampling rate of 48000 Hz; downsampling factor of 3; frame length of 32 samples; processing delay of 5 frames. Before the start of each experimental session, the amplification levels were adjusted so that the insert earphone signals were 5 dB higher than the microphone signal (Chao et al., 2019).

We collected electroencephalography (EEG) data continuously throughout the experimental session using a high-density EEG system (Brainvision) with active Ag/AgCl electrodes. Before they were seated in the sound booth, participants were fitted with an appropriately sized elastic cap (actiCAP with SNAP holders). We recorded EEG signals from 64 scalp sites using the standard 10-10 electrode system. We used four passive Ag/AgCl electrodes to record electrooculography (EOG) signals (one electrode placed superior to the participant’s right eyebrow midpoint and one inferior to their medial left eye), and electromyography (EMG) signals (one each on the left and right inferior orbicularis oris). We also used an additional microphone (Shure MVL Lavalier) to collect speech signals alongside EEG, EOG, and EMG. We used the PyCorder software, a Python-based open-source acquisition program, to collect all electrophysiological signals at a sampling rate of 1000 Hz.

### Procedure

Data collection occurred during a single 3-hour session. Following an initial training and pre-test block, participants performed 8 compensation and 4 silent reading blocks. These 12 experimental blocks were organized into four cycles, each consisting of two compensation blocks and one reading block, presented in a randomized order.

### Training Task

The practice task served to familiarize participants with the experimental protocol (∼2 minutes). Each trial began with a red target word (/hɛp/, /hɛd/, or /hɛk/; Arial Rounded MT Bold, 220 pt) centered on a gray background for 800 ms. To maintain consistency with the subsequent compensation task, an auditory probe (1 kHz, 50 ms, 75 dB SPL, 5ms rise/fall time) was delivered 500 ms after stimulus onset in 50% of the trials, occurring during the speech planning phase. Following the initial 800 ms, the target word turned green for 2,000 ms, serving as the “go” signal for the participant to produce the word aloud. Participants were instructed to initiate speech within 700 ms of the green cue and were required to maintain a target intensity of 68–78 dB SPL and a vowel duration of 500–800 ms. Visual feedback regarding speech latency (relative to the go signal), vowel intensity, and vowel duration was provided immediately following each production for 750 ms. Each trial concluded with a blank screen for a randomized inter-trial interval of 1–1.5 s.

### Pre-test Task

To establish baseline vowel configurations, participants completed a pre-test task (∼5 minutes). The trial structure mirrored the practice task; however, the target words were /hɪp/, /hɛp/, and /hæp/ (20 repetitions of each word). Additionally, consistent with the other experimental tasks, an auditory probe (1 kHz, 50 ms, 75 dB SPL, 5ms rise/fall time) was presented during the planning stage (500 ms from the onset of the red word) in 50% of the trials. In this task, we did not provide visual feedback regarding speech latency, intensity, or duration. After the task concluded, we calculated the mean formant estimates using Audapter and visually inspected the vowel distributions in the two-dimensional formant space. We then determined the centroids of the distributions of /ɪ/, /ɛ/, and /æ/. Additionally, we quantified the spatial relationship between /ɛ/ and /æ/ by calculating their relative distance and angle. These metrics were foundational for the subsequent compensation task: formants of the ɛ-centroid served as the baseline for Audapter’s formant tracking, and the ɛ–æ vector was used to determine participant-specific perturbations. Additionally, we extracted a 50-ms window from the ɛ-centroid sample to generate a participant-specific auditory probe that was used in the subsequent compensation and reading tasks.

### Compensation and Silent Reading Tasks

Following the pre-test task, participants completed 8 blocks of a compensation task (∼5 minutes) and 4 blocks of a silent reading task (∼2 minutes). The overall structure of the two tasks was similar, except that the compensation task required participants to produce speech. Similar to the training task, each trial started with a presentation of a monosyllabic target word containing /ɛ/ (/hɛp/, /hɛd/, and /hɛk/) in red. After 800 ms, the target word changed to green; this color change served as a cue for participants to produce the word aloud in the compensation task. In 40% of trials (*perturbed trials*), participants received altered auditory feedback such that the formants of /ɛ/ were shifted toward /æ/ in the two-dimensional formant space, and no auditory perturbations were applied in the remaining 60% of trials (*unperturbed trials*). The perturbed were randomized such that there was at least one unperturbed trial before each perturbed trial. In half of the perturbed trials and two-thirds of unperturbed trials (*probe trials*), we played the participant-specific auditory probe 500 ms after the presentation of the red word (/ɛ/, 50 ms, 75 dB SPL, 5 ms rise/fall time). Therefore, the compensation tasks comprised 4 types of trials: probe trials with perturbation, probe trials without perturbation, no-probe trials with perturbation, and no-probe trials without perturbation. Because participants did not produce speech during the reading task (and thus no auditory perturbations were applied), the task included two types of trials: probe trials and no-probe trials.

### Data Analysis

#### EEG Data Analysis

Processing of the EEG signal was performed using the EEGLAB toolbox in MATLAB (Delorme & Makeig, 2004). The preprocessing of the EEG signals included several steps. First, we merged all blocks of the compensation and reading tasks for each subject into a single continuous dataset. Second, the continuous EEG signals were digitally filtered using a zero-phase Hamming windowed sinc FIR filter (as implemented in the EEGLAB *pop_eegfiltnew* function). We applied a band-pass filter with a 0.5 Hz high-pass cutoff and a 55 Hz low-pass cutoff. The software automatically determined the filter order to ensure a transition bandwidth of at least 2 Hz. Third, we performed an automated artifact rejection and channel cleaning (as implemented in the EEGLAB *pop_clean_rawdata* function). The software identified and removed bad channels if (1) channels remained flat for more than 5 s, (2) channels correlated lower than 0.85 with their reconstructed neighboring channels, and (3) channels exceeded 4 standard deviations of line-noise relative to the signal. Following channel removal, the software applied Artifact Subspace Reconstruction (ASR; (Mullen et al., 2015)) to the continuous data to correct non-stationary, high-amplitude artifacts (e.g., eye blinks, muscle activity, or movement). We used a Burst Criterion of 12 (Standard Deviations) to identify artifactual subspaces relative to a clean, automatically determined reference segment. The software used a spherical interpolation matrix based on the clean reference data to reconstruct high-variance components. Additionally, we set the BurstRejection parameter to “off” so that artifactual segments were repaired rather than deleted, thus maintaining the temporal continuity of the data for subsequent analyses. Fourth, to maintain a consistent electrode montage across all participants and to facilitate group-level analysis, previously removed channels were reconstructed using spherical spline interpolation (Perrin et al., 1989). This procedure ensured that the final dataset for each participant included all 64 electrodes, enabling standardized topographical mapping and re-referencing.

Fifth, we used an Independent Component Analysis (ICA) (as implemented in the EEGLAB *pop_runica* function) to isolate and remove stereotypical artifacts (e.g., eye blinks, eye movements, and muscle activity). For this purpose, we first created a copy of the EEG data and used that to determine ICA weights. To optimize the decomposition, we down-sampled the EEG signals to 250 Hz and high-pass filtered them at 1.0 Hz (using a zero-phase FIR filter), as this approach improves ICA stability and decomposition quality (Winkler et al., 2015). Because previous preprocessing steps reduced the data’s rank, we used a Principal Component Analysis (PCA) to reduce the dimensionality based on the number of interpolated channels. This approach prevented rank deficiency and ensured stable ICA convergence. We then ran the ICA algorithm with a learning rate of 10^−5^ for up to 2,000 iterations to ensure precise source separation. After convergence, the resulting ICA weights were applied to the original dataset to reject artifacts. Following the ICA decomposition, we automatically categorized independent components (as implemented in EEGLAB ICLabel function) (Pion-Tonachini et al., 2019). This software uses a pre-trained model to assign each component a probability score across seven categories: Brain, Muscle, Eye, Heart, Line Noise, Channel Noise, and Other. This automated approach ensured objective, standardized identification of artifacts, eliminating the potential bias associated with manual component inspection. We pruned the components based on the probability scores generated by ICLabel. Specifically, any component classified as Eye, Muscle, Line Noise, or Channel Noise with a probability exceeding 80% was flagged for removal. These artifactual components were then subtracted from the data. By removing these non-brain components, we corrected eye and muscle artifacts while preserving the underlying neural signal for subsequent analysis.

Sixth, we re-referenced the cleaned EEG signals to the average mastoid electrodes (TP9 and TP10). We then segmented the continuous data into epochs time-locked to the onset of the auditory probes. For trials in which no auditory probe was presented, we used “pseudo-triggers” at the identical temporal offset as the auditory probes (500 ms after the red word). Each epoch spanned from -100 ms to +600 ms relative to the probe (or pseudo-probe) onset. We performed baseline correction by subtracting the mean amplitude of the 100-ms pre-stimulus interval from the entire epoch. We calculated the event-related potentials (ERPs) by averaging the epochs across trials for each condition. This procedure resulted in four distinct ERPs in the compensation task (probe with perturbation, probe without perturbations, no-probe with perturbation, and no-probe without perturbations) and two distinct ERPs in the reading task (probe and no-probe).

Finally, to isolate the neural response specific to auditory processing, we employed a subtraction method to remove task-related activity that was unrelated to the probe. The “probe” trials contained a mixture of auditory-evoked potentials and task-related activity (e.g., motor, cognitive, and visual processing), whereas the “no-probe” trials captured only the latter. By subtracting the no-probe ERPs from the probe ERPs in each task, we derived a difference waveform that represents the auditory activity of interest (Luck, 2014). This resultant signal was subsequently low-pass filtered at 30 Hz (using a zero-phase FIR filter) to generate the final ERPs for analysis, a procedure consistent with established protocols for controlling overlapping neural activity (Baess et al., 2011; Daliri & Max, 2015b; Klaffehn et al., 2019; Luck, 2014). This procedure generated two ERPs in the compensation task (one for perturbed trials and one for unperturbed trials) and one ERP in the silent reading task. Based on prior findings indicating that pre-speech modulation of the N1 and P2 components is most prominent over frontocentral scalp sites (Daliri & Max, 2015b), we defined a Region of Interest (ROI) consisting of 20 channels (electrodes are highlighted in topographical maps in Figures 1 and 2). For each participant and condition, we averaged ERP waveforms across these 20 channels to create a single representative ROI signal. From this averaged waveform, we identified the N1 component as the maximum negative deflection occurring within a 70–130 ms post-stimulus interval, and the P2 component as the maximum positive deflection within the 150–250 ms post-stimulus interval (Kappenman & Luck, 2011; Luck, 2014; Näätänen et al., 1987). We extracted the peak amplitude and latency of the N1 and P2 components as dependent measures for statistical analyses. We defined pre-speech auditory modulation for the N1 and P2 components as the difference between their amplitudes in the compensation and reading tasks.

**Figure 1.**
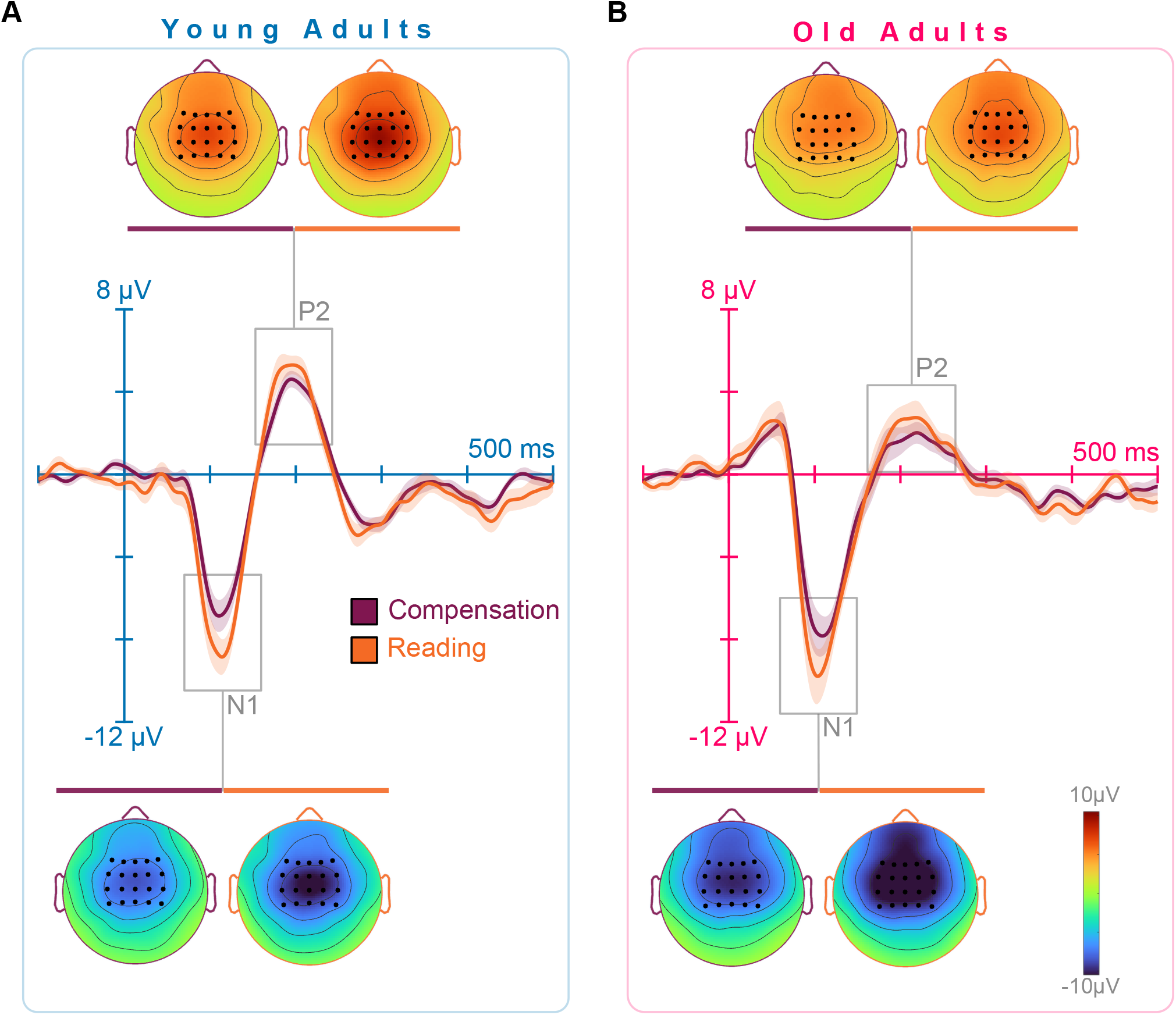
Grand average ERP waveforms and topographical distributions. Waveforms represent responses to the probe stimulus during the compensation task and silent reading for (A) young adults (n=17) and (B) old adults (n=13). Data are averaged across 20 frontocentral electrodes, indicated by dark circles on the topographic maps. Topographic maps illustrate the scalp voltage distribution for the N1 and P2 components across both tasks. Shaded areas denote the standard error of the mean.

**Figure 2.**
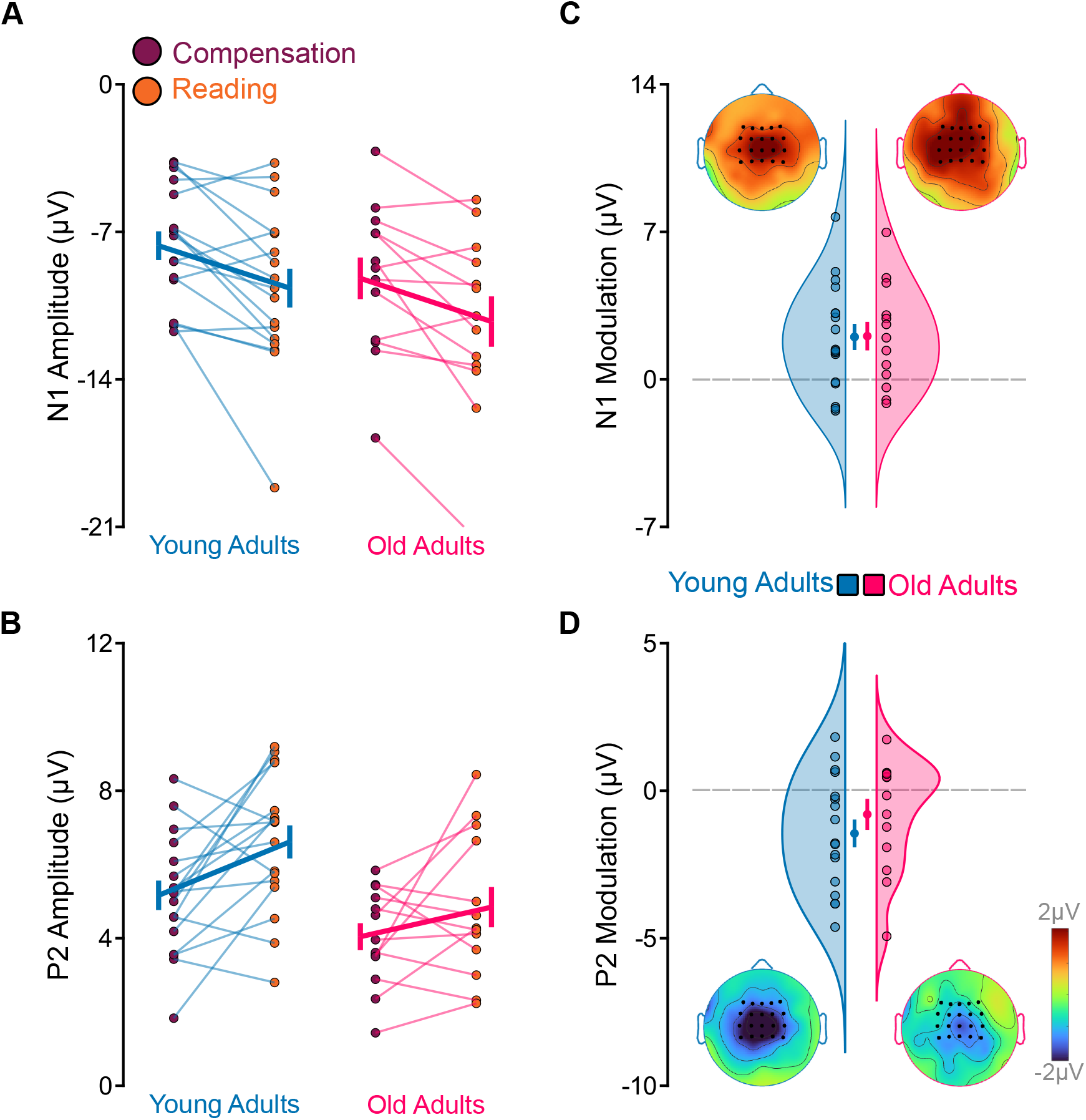
N1 and P2 amplitudes and pre-speech auditory modulation (PSAM) across age groups. (A, B) Individual data points and group means for N1 and P2 amplitudes are extracted from the frontocentral electrode cluster. (C, D) PSAM is defined as the amplitude difference between the compensation and silent reading tasks (speaking minus reading) for the N1 and P2 components. Topographical maps illustrate the scalp distribution of N1 and P2 PSAMs. Error bars represent the standard error of the mean.

**Figure 3.**
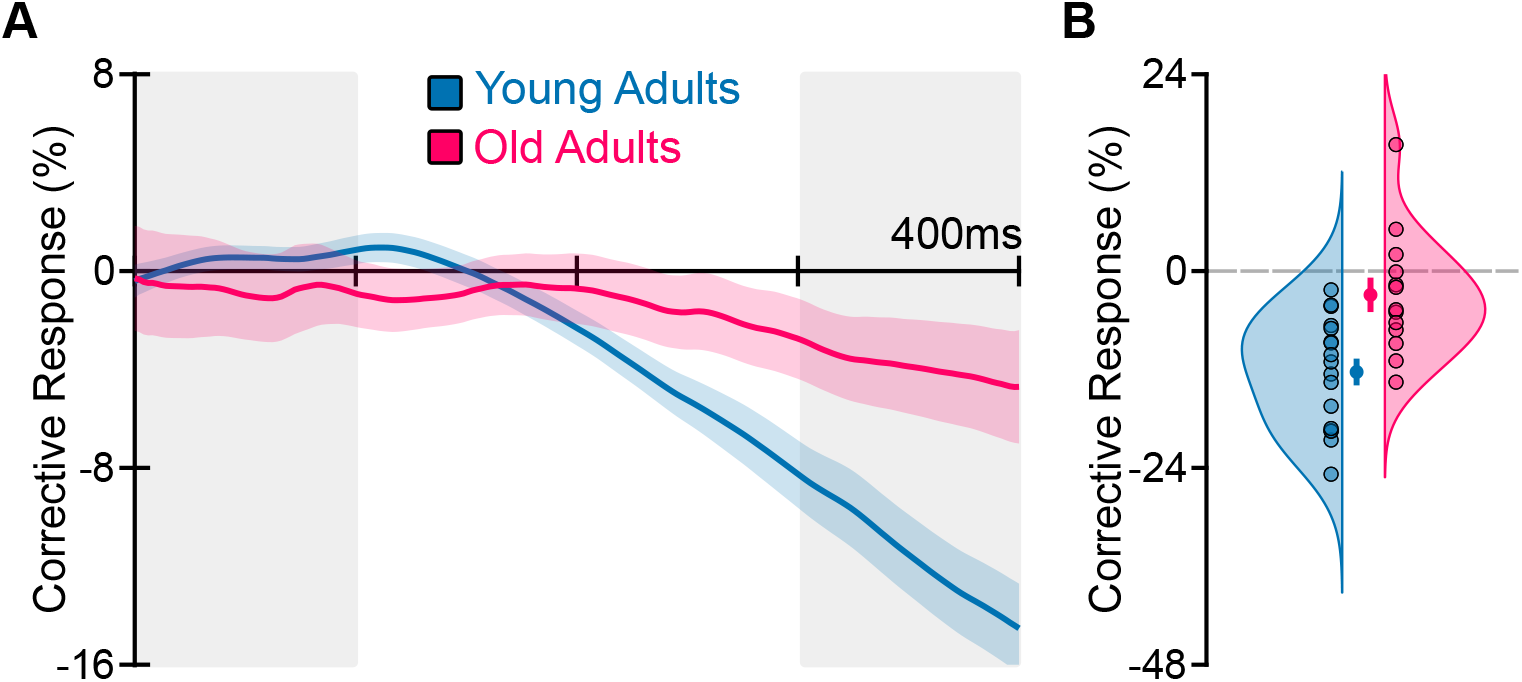
Formant trajectories and corrective responses to auditory perturbations. (A) Trajectories of formant change in F1-F2 space, projected along the perturbation vector. (B) Individual data points and group mean for corrective responses in young and old adults. Corrective responses were calculated as the difference in formant change between the final 100 ms and the initial 100 ms of the response (grey shaded areas). Old adults exhibited significantly smaller corrective responses than young adults (p = .001). Error bars represent the standard error of the mean.

**Figure 4.**
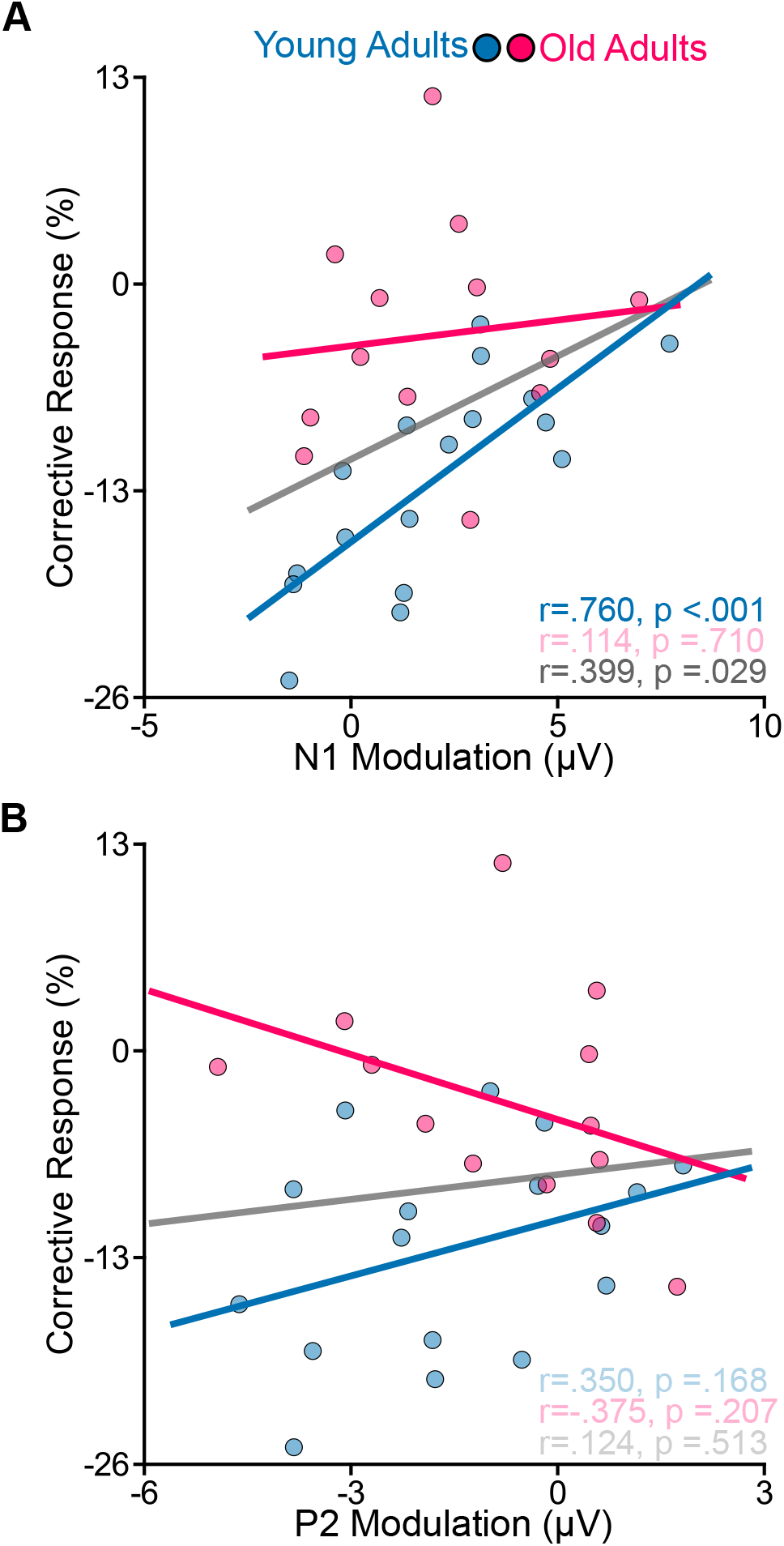
Correlations between pre-speech auditory modulation and corrective responses. (A) Relationship between N1 pre-speech auditory modulation (PSAM) and corrective responses for young and older adults. In young adults, a significant correlation was observed: larger N1 PSAM was associated with smaller corrective responses (p < .001); no such relationship was found in older adults. (B) Relationship between P2 PSAM and corrective responses. No significant correlations were observed for either age group. Scatterplots include individual data points and linear regression lines for each group.

#### Speech Data Analysis

We analyzed the acoustic data from the compensation task using a multi-stage procedure. First, an experimenter—blinded to the perturbation conditions—visually inspected all trial spectrograms and excluded productions containing mispronunciations or significant formant-tracking errors. The experimenter also manually marked the vowel onsets and offsets for each trial to define the temporal boundaries for formant extraction. To isolate the compensatory effects of the auditory perturbations, we developed participant- and word-specific formant templates using unperturbed trials. We excluded any unperturbed trials immediately following a perturbed trial from these templates to account for potential carryover effects and ensure we captured baseline speech (Sreedhar & Daliri, 2026). For each word, we resampled the baseline formant trajectories to 100 points and averaged across the selected unperturbed trials. We then interpolated these time-normalized templates to match the duration of each experimental trial and subtracted them from the raw trajectories, effectively removing word-specific formant transitions (Chao & Daliri, 2023; Daliri et al., 2020; Merrikhi et al., 2025; Sreedhar & Daliri, 2026). We extracted formants from the first 400 ms of the vowel and excluded trials shorter than this duration. Overall, we excluded 3.4% (SD = 6.2%) of all trials across participants. Consistent with our previous work, we projected the F1 and F2 responses onto a single axis in the F1−F2 coordinate system, such that negative values indicated a change in the opposite direction of the ε−æ perturbation. We defined the corrective response as the within-trial change in formants, calculated by subtracting the mean trajectory of the 0–100 ms window from that of the 300–400 ms window. We used these corrective responses as the dependent measures for all subsequent statistical analyses.

#### Statistical Analysis

We conducted all statistical analyses using R version 4.3.3 (R. C. Team, 2021) in the RStudio environment version 2023.12.1 (Rs. Team, 2020). Our original design for the compensation task included four trial types: (1) no-probe/unperturbed, (2) no-probe/perturbed, (3) probe/unperturbed, and (4) probe/perturbed. During ERP preprocessing, we isolated auditory activity by subtracting no-probe averages from probe averages. Because our preliminary analysis revealed no significant differences in ERPs between perturbed and unperturbed trials, we report only the perturbed-trial ERPs, which directly address our primary research questions. Similarly, we calculated corrective responses by subtracting the average of unperturbed trials from the perturbed trials for both probe and no-probe trials. Since the presence of an auditory probe during the planning stage did not significantly affect the magnitude of corrective responses, we do not report results for no-probe trials. These simplifications streamlined the statistical analysis while preserving the focus on the critical experimental contrasts.

We used the lme4 package (version 1.1.35.1) to fit a linear mixed-effects model for each dependent variable (Bates et al., 2015). For models fitted to dependent measures derived from the ERPs, we used group (young adults vs. old adults) and task (compensation vs. reading) as fixed effects. For models fitted with the corrective responses, we used group (young adults vs. old adults) as a fixed effect. We first sought to include a maximal random-effects structure, including random slopes for the interaction between group and condition. However, this complex specification resulted in singular fits or non-convergence, even after we followed a systematic reduction approach (Barr et al., 2013). To ensure model stability and avoid over-parameterization, we ultimately used a random-intercept-only structure, which accounted for baseline inter-participant variability. We verified the model assumptions—specifically the normality of residuals and homoscedasticity—by inspecting diagnostic plots (Harrison et al., 2018). We determined the significance of fixed effects using the lmerTest package (version 3.1.3), employing Satterthwaite’s approximation for estimating degrees of freedom (Kuznetsova et al., 2017). For further analysis of significant fixed effects and interactions, we used the emmeans package (version 1.8.7) to estimate marginal means and conduct post hoc contrasts (Lenth, 2019). We applied the Kenward-Roger method to adjust the degrees of freedom and standard errors for these comparisons and controlled the family-wise error rate using False Discovery Rate (FDR) corrections. To examine the relationships between behavioral and neural measures, we calculated Pearson correlation coefficients using the psych package (version 2.4.1; (Revelle, 2018)). We then compared these correlations between groups using a nonparametric permutation test (10,000 iterations). For this procedure, we first calculated the actual difference between group correlation coefficients. We then generated a null distribution by pooling the data and randomly reassigning group labels while keeping each participant’s neural and behavioral data pairs intact. We defined the p-value as the proportion of permutations that yielded a difference equal to or more extreme than our observed result (Zar, 2010). This permutation-based approach enabled us to evaluate group differences in correlation strength without assuming bivariate normality.

## Results

### Pre-Speech Auditory Modulation (PSAM) and Corrective Responses

Figure I shows the grand average ERP waveforms elicited by the probe stimulus during the compensation and silent reading tasks for both young and older adults. From these waveforms, we extracted peak amplitudes and peak latencies for the N1 and P2 components. To facilitate statistical analysis, we averaged the data across a cluster of 20 frontocentral electrodes, which are highlighted in the topographical maps in Figure I. Figure I also illustrates the topographical distributions of N1 and P2 for both age groups and task conditions. Examining N1 amplitude, we found a statistically significant main effect of task, F(1, 28) = 19.345, p < .001, indicating that N1 amplitude was smaller in the compensation task than in the reading task (i.e., a statistically significant N1 PSAM). However, the main effect of group (p = .220) and group × task interaction (p = .966) were not statistically significant. Examining P2 amplitude, we found statistically significant main effects of task, F(1, 28) = 10.076, p = .004, and group, F(1, 28) = 7.605, p = .010. However, the group × task interaction was not statistically significant (p = .369). These findings indicated that participants showed significant P2 PSAM and that the overall P2 amplitude in young adults was larger than that in old adults. Figure II A and II B present the individual data points and grand-average amplitudes for the N1 and P2 components across both tasks. In panels C and D, we illustrate the N1 and P2 PSAMs for both age groups, accompanied by topographical maps detailing the scalp distribution of these modulations.

Examining N1 latency, we did not find a statistically significant main effect of task (p = .441), a main effect of group (p = .129), or a group × task interaction (p = .608). The average N1 latency for young adults was 112 ± 9.6 ms (mean ± SD), and for old adults, 106 ± 13.3 ms. Examining P2 latency, we found a statistically significant main effect of group, F(1,28) = 7.712, p = .010, indicating that the average P2 latency of young adults (196 ± 14.7 ms) was shorter than that of old adults (215 ± 27.0 ms). We did not find a statistically significant main effect of task (p = .660) or group × task interaction (p = .156).

Finally, our analysis of the corrective responses indicated that both groups produced corrective responses to the perturbations; however, young adults produced larger corrective responses than old adults (t(25.144) = -3.671, p = .001). Figure III A illustrates the temporal trajectories of corrective responses, while Figure III B presents the individual data points alongside the grand averages for both age groups.

### Relationship between Pre-Speech Auditory Modulation and Corrective Responses

The primary objective of this study was to examine the relationship between PSAM and corrective responses to auditory errors. As shown in Figure IV A, N1 PSAM was correlated with corrective responses in young adults (r = .760, p < .001), but not in old adults (r = .114, p = .710). Our nonparametric analysis showed that the correlation coefficient for young adults differed significantly from that for old adults (p = .004). This analysis indicated that young adults who modulated their auditory system more during speech planning also responded less to auditory errors they experienced during speech production. However, the relationship between P2 PSAM and corrective responses (Figure IV B) was not significant in either young adults (r = .350, p = .168) or old adults (r = -.375, p = .207). No dependent variables were significantly correlated with chronological age or MMSE scores (p > .350).

## Discussion

Studies have shown that the brain modulates the auditory system as it prepares speech movements (Daliri & Max, 2015b; Eliades & Wang, 2008; Seki & Fetz, 2012). We recently proposed that pre-speech auditory modulation (PSAM) serves as a neural index of the predictive processes underlying feedforward motor preparation (Li et al., 2024). Predictive processes also play an essential role in error detection and correction during production (Sreedhar & Daliri, 2026; Tourville et al., 2008). Therefore, the primary objective of this study was to elucidate the relationship between PSAM and corrective responses to auditory errors. Furthermore, a secondary objective of this study was to characterize PSAM and corrective responses in both young and old adults to determine the impact of aging on predictive processes. To address these objectives, we measured PSAM by comparing neural activity during speaking and silent reading conditions in young and old adults. We also measured corrected responses to perturbations that shifted the formants of /ɛ/ toward /æ/. Our results revealed that (1) while N1 PSAM and P2 PSAM were significant for both young and old adults, only N1 PSAM of young adults correlated with their corrective responses to auditory errors, and (2) older adults had smaller and delayed P2 components along with smaller corrective responses to the auditory errors.

Our finding of a significant correlation between N1 PSAM and corrective responses in young adults supports the hypothesis that PSAM serves as a neural index of the brain’s reliance on predictive processes (with aging as a confound). Specifically, we observed that young individuals with larger N1 PSAMs exhibited reduced responses to auditory errors induced by speech perturbations. The brain relies on a balance between internal predictions and sensory feedback (Daliri, 2021; Guenther, 2016; Houde & Nagarajan, 2011; Parrell & Houde, 2019). When the brain places greater weight on its own predictions, it may “discount” feedback-driven error signals, resulting in reduced corrective responses. Thus, the increased reliance on prediction, as indexed by a larger PSAM, likely explains the reduced corrective responses observed here. These results also allow us to reassess competing interpretations of PSAM. Previously, we and others suggested that PSAM might reflect the “fine-tuning” of the auditory system to enhance monitoring accuracy (Eliades & Wang, 2008, 2017; Max & Daliri, 2019; Schneider et al., 2014). However, both our previous work and the current study provide evidence against this interpretation. In our previous study (Li et al., 2024), PSAM correlated with trial-to-trial speech variability at vowel onset—a measure of feedforward control accuracy. If PSAM truly served as a fine-tuning mechanism for more accurate error detection, one would expect larger PSAM to correlate with greater sensitivity and, consequently, larger corrective responses. The current study’s findings are opposite to this prediction—larger modulation is associated with smaller corrective responses. This inverse relationship contradicts the fine-tuning hypothesis. Collectively, our findings regarding trial-to-trial variability and corrective responses reinforce the theory that PSAM is a marker of predictive reliance, playing a fundamental role in both feedforward and feedback control mechanisms.

The second objective of this study was to evaluate the impact of aging on predictive processes in speech motor control. The older adults as a group exhibited a qualitative increase in their auditory P50 ERP in response to the probe tones, consistent with previously reported age-related changes in sensory gating (Stothart & Kazanina, 2016). However, despite these broader age-related changes, N1 PSAM itself was not significantly impacted by age, though older adults had smaller corrective responses. Thus, aging seemed to significantly decouple the relationship between this neural index and behavioral corrective responses. While these findings together suggest predictive processing may be preserved in aging, they do not specify how predictive and corrective cues may be differently weighed in older adults. Reduced corrective responses in older adults could be explained by several possible mechanisms affecting speech motor control. Perhaps aging impairs the processes underlying error detection and correction rather than the internal predictive mechanisms themselves. Due to their age-related decline in the neural substrates of sensory processing (Bornkessel-Schlesewsky et al., 2015; Ferreri et al., 2017) older adults may place less weight on sensory feedback or experience an altered balance between prediction and feedback, leading them to discount auditory errors (Chan et al., 2021; Moran et al., 2014). Increase in sensory noise could also cause older adults to rely more on predictive processing in state estimation, ultimately leading to decreased responses to sensory errors (de Witte et al., 2026; Wolpe et al., 2016). However, this scenario is less likely as the older adult group in the current study did not have significantly more modulation compared to young adults. Alternatively, aging may impair the ability to convert perceived errors into motor corrections.

Because error correction relies heavily on frontal lobe motor regions (for a review, see Guenther, 2016)—areas particularly vulnerable to age-related atrophy (C. Clark & L. Taylor, 2011; Lemaitre et al., 2012; Storsve et al., 2014)—the motor execution of a correction may be compromised even if the error is correctly identified. Aging also significantly influenced the P2 component, resulting in smaller amplitudes and increased latencies. Given that P2 reflects higher-stage sensory processing and has neural generators distributed across the age-sensitive frontal and temporal lobes (Kappenman & Luck, 2011; Luck, 2014; Näätänen et al., 1987), these changes are consistent with broad age-related neurodegeneration (C. Clark & L. Taylor, 2011; Ferreri et al., 2017; Fjell et al., 2014; Lemaitre et al., 2012; Storsve et al., 2014). Crucially, however, the P2 PSAM remained unaffected by age. This stability indicates that the specific predictive processes indexed by P2 PSAM are likely resilient to the effects of aging. Overall, N1 PSAM appeared relatively preserved in older adults despite reduced corrective responses to auditory perturbation. Multiple mechanisms may account for this dissociation, and further work is needed to draw definitive mechanistic conclusions regarding speech motor control in aging populations. Despite the insights this study provides, several limitations warrant consideration. First, the relationship between N1 PSAM during planning and corrective responses during production remains correlational. Although PSAM temporally precedes the corrective response, and our measurements were taken from the same trials to ensure maximum temporal proximity, these findings do not confirm a causal link. Future research should employ neuromodulation techniques (e.g., transcranial magnetic stimulation) to directly manipulate predictive processes and observe their downstream effects on corrective responses. Second, because our analyses were restricted to scalp-level EEG data, our ability to draw conclusions regarding the specific neural generators of these signals is limited. Incorporating source localization into future studies will be essential to precisely identify the cortical regions involved and clarify how aging affects these generators. Finally, while this study compared young and old adults, a full understanding of these mechanisms requires a lifespan perspective. Future studies should include pediatric populations to characterize how the balance between predictive and feedback-driven control develops from childhood through late adulthood.

In summary, the primary objective of the current study was to elucidate the relationship between pre-speech auditory modulation and corrective responses to auditory errors. We measured auditory modulation by comparing neural activity during speaking and silent reading conditions in young and old adults. While speaking, participants also experienced auditory perturbations (/ɛ/ sounded like /æ/). Our results revealed that (1) while N1 modulation and P2 modulation were significant for both young and old adults, only N1 modulation of young adults correlated with their corrective responses to auditory errors, and (2) old adults had smaller and delayed P2 components along with smaller corrective responses to the auditory errors. Overall, these findings reinforce the theory that pre-speech auditory modulation is a marker of reliance on predictive processes, which play a fundamental role in both feedforward and feedback control mechanisms. Additionally, our findings also suggest that aging impairs the processes underlying error detection and correction rather than the internal predictive mechanisms themselves.

## Acknowledgments

This work was supported by a grant from the National Institutes of Health awarded to A. Daliri (R01-DC019905 and R01-DC020162). The content is solely the responsibility of the authors and does not necessarily represent the official views of the National Institutes of Health.

The manuscript’s grammar and syntax were refined using the Gemini 1.5 Pro large language model (Google). Following the AI-assisted editing, the authors critically reviewed and revised all output to ensure technical accuracy and stylistic consistency before final inclusion.

## Data Availability Statement

The dataset collected for the current study is available upon reasonable request from the second author (A. Daliri).

## Notes

### Competing Interest Statement

The authors have declared no competing interest.

## References

ASHA. (1997). Guidelines for audiologic screening: Panel on audiologic assessment. American Speech-Language-Hearing Association.

Baess, P., Horváth, J., Jacobsen, T., & Schröger, E. (2011). Selective suppression of self-initiated sounds in an auditory stream: An ERP study. Psychophysiology, 48(9), 1276–1283. 10.1111/j.1469-8986.2011.01196.x

Barr, D. J., Levy, R., Scheepers, C., & Tily, H. J. (2013). Random effects structure for confirmatory hypothesis testing: Keep it maximal. Journal of Memory and Language, 68(3), 255–278. 10.1016/j.jml.2012.11.001

Bates, D., Mächler, M., Bolker, B. M., & Walker, S. C. (2015). Fitting linear mixed-effects models using lme4. Journal of Statistical Software, 67(1), 1–48. 10.18637/jss.v067.i01.

Bays, P. M., Wolpert, D. M., & Flanagan, J. R. (2005). Perception of the consequences of self-action is temporally tuned and event driven. Current Biology, 15(12), 1125–1128. 10.1016/j.cub.2005.05.023

Behroozmand, R., Khoshhal Mollasaraei, Z., Nejati, V., Daliri, A., & Fridriksson, J. (2025). Vocal and articulatory speech control deficits in individuals with post-stroke aphasia. Scientific Reports, 15(1), 13350. 10.1038/s41598-025-96040-4

Blakemore, S.-J., Wolpert, D. M., & Frith, C. D. (1998). Central cancellation of self-produced tickle sensation. Nature Neuroscience, 1(7), 635–640. 10.1038/2870

Bornkessel-Schlesewsky, I., Philipp, M., Alday, P. M., Kretzschmar, F., Grewe, T., Gumpert, M., Schumacher, P. B., & Schlesewsky, M. (2015). Age-Related Changes in Predictive Capacity Versus Internal Model Adaptability: Electrophysiological Evidence that Individual Differences Outweigh Effects of Age. Frontiers in Aging Neuroscience, 7. 10.3389/fnagi.2015.00217

C. Clark, B., & L. Taylor, J. (2011). Age-Related Changes in Motor Cortical Properties and Voluntary Activation of Skeletal Muscle. Current Aging Sciencee, 4(3), 192–199. 10.2174/1874609811104030192

Cai, S. (2015). Audapter. https://github.com/shanqing-cai/audapter_matlab

Cai, S., Beal, D. S., Ghosh, S. S., Tiede, M. K., Guenther, F. H., & Perkell, J. S. (2012). Weak responses to auditory feedback perturbation during articulation in persons who stutter: Evidence for abnormal auditory-motor transformation. PLoS ONE, 7(7), e41830. 10.1371/journal.pone.0041830

Chan, J. S., Wibral, M., Stawowsky, C., Brandl, M., Helbling, S., Naumer, M. J., Kaiser, J., & Wollstadt, P. (2021). Predictive Coding Over the Lifespan: Increased Reliance on Perceptual Priors in Older Adults—A Magnetoencephalography and Dynamic Causal Modeling Study. Frontiers in Aging Neuroscience, 13, 631599. 10.3389/fnagi.2021.631599

Chang, S.-E., & Guenther, F. H. (2020). Involvement of the cortico-basal ganglia-thalamocortical loop in developmental stuttering. Frontiers in Psychology, 10(3088). 10.3389/fpsyg.2019.03088

Chao, S., & Daliri, A. (2023). Effects of gradual and sudden introduction of perturbations on adaptive responses to formant-shift and formant-clamp perturbations. Journal of Speech, Language, and Hearing Research, 66(5), 1588–1599. 10.1044/2023_JSLHR-21-00435

Chao, S., Ochoa, D., & Daliri, A. (2019). Production variability and categorical perception of vowels are strongly linked. Frontiers in Human Neuroscience, 13, 96. 10.3389/FNHUM.2019.00096

Daliri, A. (2021). A computational model for estimating the speech motor system’s sensitivity to auditory prediction errors. Journal of Speech, Language, and Hearing Research, 64(6), 1841–1854. 10.1044/2021_JSLHR-20-00484

Daliri, A., Chao, S., & Fitzgerald, L. C. (2020). Compensatory responses to formant perturbations proportionally decrease as perturbations increase. Journal of Speech, Language, and Hearing Research, 63(10), 3392–3407. 10.1044/2020_JSLHR-19-00422

Daliri, A., & Dittman, J. (2019). Successful auditory motor adaptation requires task-relevant auditory errors. Journal of Neurophysiology, 122(2), 552–562. 10.1152/jn.00662.2018

Daliri, A., Honda, S., & Max, L. (2025). Delayed auditory feedback increases speech production variability in typically fluent adults but has the opposite effect in stuttering adults. Frontiers in Human Neuroscience, 19, 1628114. 10.3389/fnhum.2025.1628114

Daliri, A., & Max, L. (2015a). Electrophysiological evidence for a general auditory prediction deficit in adults who stutter. Brain and Language, 150, 37–44. 10.1016/j.bandl.2015.08.008

Daliri, A., & Max, L. (2015b). Modulation of auditory processing during speech movement planning is limited in adults who stutter. Brain and Language, 143, 59–68. 10.1016/j.bandl.2015.03.002

Daliri, A., & Max, L. (2016). Modulation of auditory responses to speech vs. Nonspeech stimuli during speech movement planning. Frontiers in Human Neuroscience, 10, 1–9. 10.3389/fnhum.2016.00234

Daliri, A., & Max, L. (2018). Stuttering adults’ lack of pre-speech auditory modulation normalizes when speaking with delayed auditory feedback. Cortex, 99, 55–68. 10.1016/j.cortex.2017.10.019

Daliri, A., Prokopenko, R. A., Flanagan, J. R., & Max, L. (2014). Control and prediction components of movement planning in stuttering versus nonstuttering adults. Journal of Speech, Language, and Hearing Research, 57(6), 2131–2141. 10.1044/2014_JSLHR-S-13-0333

de Witte, A., Matthijs, A., Parrell, B., Mantini, D., Gooijers, J., & Orban de Xivry, J. J. (2025). Preserved cerebellar functions despite structural degeneration in older adults. bioRxiv, 2025-06.

Delorme, A., & Makeig, S. (2004). EEGLAB: An open source toolbox for analysis of single-trial EEG dynamics including independent component analysis. Journal of Neuroscience Methods, 134(1), 9–21. 10.1016/j.jneumeth.2003.10.009

Eliades, S. J., & Wang, X. (2008). Neural substrates of vocalization feedback monitoring in primate auditory cortex. Nature, 453(7198), 1102–1106. 10.1038/nature06910

Eliades, S. J., & Wang, X. (2017). Contributions of sensory tuning to auditory-vocal interactions in marmoset auditory cortex. Hearing Research, 348, 98–111. 10.1016/j.heares.2017.03.001

Ferreri, F., Guerra, A., Vollero, L., Ponzo, D., Maatta, S., Mervaala, E., Iannello, G., & Lazzaro, V. D. (2017). Age-related changes of cortical excitability and connectivity in healthy humans: Non-invasive evaluation of sensorimotor network by means of TMS-EEG. Neuroscience, 357, 255–263. 10.1016/j.neuroscience.2017.06.014

Fjell, A. M., Westlye, L. T., Grydeland, H., Amlien, I., Espeseth, T., Reinvang, I., Raz, N., Dale, A. M., Walhovd, K. B., & for the Alzheimer Disease Neuroimaging Initiative. (2014). Accelerating Cortical Thinning: Unique to Dementia or Universal in Aging? Cerebral Cortex, 24(4), 919–934. 10.1093/cercor/bhs379

Folstein, M. F., Folstein, S. E., & McHugh, P. R. (1975). “Mini-mental state.” Journal of Psychiatric Research, 12(3), 189–198. 10.1016/0022-3956(75)90026-6

Frankford, S. A., Cai, S., Nieto-Castañón, A., & Guenther, F. H. (2022). Auditory feedback control in adults who stutter during metronome-paced speech II. Formant Perturbation. Journal of Fluency Disorders, 74, 105928. 10.1016/J.JFLUDIS.2022.105928

Guenther, F. H. (2016). Neural control of speech. MIT Press.

Harrison, X. A., Donaldson, L., Correa-Cano, M. E., Evans, J., Fisher, D. N., Goodwin, C. E. D., Robinson, B. S., Hodgson, D. J., & Inger, R. (2018). A brief introduction to mixed effects modelling and multi-model inference in ecology. PeerJ, 6, e4794. 10.7717/peerj.4794

Houde, J. F., & Jordan, M. I. (1998). Sensorimotor adaptation in speech production. Science, 279(5354), 1213–1216. 10.1126/science.279.5354.1213

Houde, J. F., & Nagarajan, S. S. (2011). Speech production as state feedback control. Frontiers in Human Neuroscience, 5, 82. 10.3389/fnhum.2011.00082

Hughes, G., Desantis, A., & Waszak, F. (2013). Mechanisms of intentional binding and sensory attenuation: The role of temporal prediction, temporal control, identity prediction, and motor prediction. Psychological Bulletin, 139(1), 133–151. 10.1037/a0028566

Jackson, E. S., Tiede, M., Beal, D., & Whalen, D. H. (2016). The impact of social–cognitive stress on speech variability, determinism, and stability in adults who do and do not stutter. *Journal of Speech*, Language, and Hearing Research, 59(6), 1295–1314. 10.1044/2016_JSLHR-S-16-0145

Kappenman, E. S., & Luck, S. J. (Eds.). (2011). The Oxford Handbook of Event-Related Potential Components (1st ed.). Oxford University Press. 10.1093/oxfordhb/9780195374148.001.0001

Klaffehn, A. L., Baess, P., Kunde, W., & Pfister, R. (2019). Sensory attenuation prevails when controlling for temporal predictability of self- and externally generated tones. Neuropsychologia, 132(July), 107145. 10.1016/j.neuropsychologia.2019.107145

Kleiner, M., Brainard, D., Pelli, D., Ingling, A., Murray, R., & Broussard, C. (2007). What’s new in psychtoolbox-3. Perception, 36(14), 1–16.

Kuznetsova, A., Brockhoff, P. B., & Christensen, R. H. B. (2017). lmerTest package: Tests in linear mixed effects models. Journal of Statistical Software, 82(13), 1–26. doi:%2010.18637/jss.v082.i13

Lametti, D. R., Rochet-Capellan, A., Neufeld, E., Shiller, D. M., & Ostry, D. J. (2014). Plasticity in the human speech motor system drives changes in speech perception. Journal of Neuroscience, 34(31), 10339–10346. 10.1523/JNEUROSCI.0108-14.2014

Lemaitre, H., Goldman, A. L., Sambataro, F., Verchinski, B. A., Meyer-Lindenberg, A., Weinberger, D. R., & Mattay, V. S. (2012). Normal age-related brain morphometric changes: Nonuniformity across cortical thickness, surface area and gray matter volume? Neurobiology of Aging, 33(3), 617.e1–617.e9. 10.1016/j.neurobiolaging.2010.07.013

Lenth, R. (2019). emmeans: Estimated marginal means. R package. https://cran.r-project.org/package=emmeans

Li, J. J., Daliri, A., Kim, K. S., & Max, L. (2024). Does pre-speech auditory modulation reflect processes related to feedback monitoring or speech movement planning? Neuroscience Letters, 843, 138025. 10.1016/J.NEULET.2024.138025

Loucks, T. M., Chon, H., & Han, W. (2012). Audiovocal integration in adults who stutter. International Journal of Language and Communication Disorders, 47(4), 451–456. 10.1111/j.1460-6984.2011.00111.x

Luck, S. J. (2014). An Introduction to the Event-Related Potential Technique (second). MIT Press.

Max, L., & Daliri, A. (2019). Limited pre-speech auditory modulation in individuals who stutter: Data and hypotheses. Journal of Speech, Language, and Hearing Research, 62(8S), 3071–3084. 10.1044/2019_JSLHR-S-CSMC7-18-0358

McGuffin, B. J., Liss, J. M., & Daliri, A. (2020). The orofacial somatosensory system is modulated during speech planning and production. Journal of Speech, Language, and Hearing Research, 63(8), 2637–2648.

Merrikhi, Y., Ebrahimpour, R., & Daliri, A. (2018). Perceptual manifestations of auditory modulation during speech planning. Experimental Brain Research, 236(7), 1963–1969. 10.1007/s00221-018-5278-3

Merrikhi, Y., Parsa, M., & Daliri, A. (2025). An integrated approach to concurrently measure corrective and adaptive responses to auditory errors. Journal of Speech, Language, and Hearing Research, 1–11. 10.1044/2025_jslhr-24-00572

Moran, R. J., Symmonds, M., Dolan, R. J., & Friston, K. J. (2014). The Brain Ages Optimally to Model Its Environment: Evidence from Sensory Learning over the Adult Lifespan. PLoS Computational Biology, 10(1), e1003422. 10.1371/journal.pcbi.1003422

Mullen, T. R., Kothe, C. A. E., Chi, Y. M., Ojeda, A., Kerth, T., Makeig, S., Jung, T.-P., & Cauwenberghs, G. (2015). Real-time neuroimaging and cognitive monitoring using wearable dry EEG. IEEE Transactions on Biomedical Engineering, 62(11), 2553–2567. 10.1109/TBME.2015.2481482

Näätänen, R., Picton, T., Rrsro, N., & Prcron, T. (1987). The N1 wave of the human electric and magnetic response to sound: A review and an analysis of the component structure. Psychophysiology, 24(4), 375–425.

Neef, N. E., & Chang, S.-E. (2024). Knowns and unknowns about the neurobiology of stuttering. PLOS Biology, 22(2), e3002492. 10.1371/journal.pbio.3002492

Parrell, B., Agnew, Z. K., Nagarajan, S. S., Houde, J. F., & Ivry, R. B. (2017). Impaired feedforward control and enhanced feedback control of speech in patients with cerebellar degeneration. Journal of Neuroscience, 37(38), 3363–16. 10.1523/JNEUROSCI.3363-16.2017

Parrell, B., & Houde, J. F. (2019). Modeling the role of sensory feedback in speech motor control and learning. Journal of Speech, Language, and Hearing Research, 62, 2963–2985.

Perrin, F., Pernier, J., Bertrand, O., & Echallier, J. F. (1989). Spherical splines for scalp potential and current density mapping. Electroencephalography and Clinical Neurophysiology, 72(2), 184–187. 10.1016/0013-4694(89)90180-6

Pion-Tonachini, L., Kreutz-Delgado, K., & Makeig, S. (2019). ICLabel: An automated electroencephalographic independent component classifier, dataset, and website. NeuroImage, 198, 181–197. 10.1016/j.neuroimage.2019.05.026

Revelle, W. (2018). psych: Procedures for psychological, psychometric, and personality research. Northwestern University. https://cran.r-project.org/package=psych

Schneider, D. M., Nelson, A., & Mooney, R. (2014). A synaptic and circuit basis for corollary discharge in the auditory cortex. Nature, 513(7517), 189–194. 10.1038/nature13724

Seki, K., & Fetz, E. E. (2012). Gating of Sensory Input at Spinal and Cortical Levels during Preparation and Execution of Voluntary Movement. Journal of Neuroscience, 32(3), 890–902. 10.1523/JNEUROSCI.4958-11.2012

Shadmehr, R., & Mussa-Ivaldi, S. (2012). Biological learning and control: How the brain builds representations, predicts events, and makes decisions. MIT Press. https://mitpress.mit.edu/books/biological-learning-and-control

Smith, A., & Kleinow, J. (2000). Kinematic correlations of speaking rate changes in stuttering and normally fluent adults. *Journal of Speech*, Language, and Hearing Research, 10(2), 127–142.

Sreedhar, A. J., & Daliri, A. (2026). The complex impact of auditory error direction, magnitude, and exposure on corrective responses. Frontiers in Human Neuroscience, 19, 1729850. 10.3389/fnhum.2025.1729850

Storsve, A. B., Fjell, A. M., Tamnes, C. K., Westlye, L. T., Overbye, K., Aasland, H. W., & Walhovd, K. B. (2014). Differential Longitudinal Changes in Cortical Thickness, Surface Area and Volume across the Adult Life Span: Regions of Accelerating and Decelerating Change. Journal of Neuroscience, 34(25), 8488–8498. 10.1523/JNEUROSCI.0391-14.2014

Stothart, G., & Kazanina, N. (2016). Auditory perception in the aging brain: the role of inhibition and facilitation in early processing. Neurobiology of aging, 47, 23–34.

Team, R. C. (2021). R: A language and environment for statistical computing. R Foundation for Statistical Computing. https://www.r-project.org/

Team, Rs. (2020). RStudio: Integrated Development for R. RStudio, PBC.

Tourville, J. A., Reilly, K. J., & Guenther, F. H. (2008). Neural mechanisms underlying auditory feedback control of speech. NeuroImage, 39(3), 1429–1443. 10.1016/j.neuroimage.2007.09.054

Villacorta, V. M., Perkell, J. S., & Guenther, F. H. (2007). Sensorimotor adaptation to feedback perturbations of vowel acoustics and its relation to perception. The Journal of the Acoustical Society of America, 122(4), 2306–2319. 10.1121/1.2773966

Waszak, F., Cardoso-Leite, P., & Hughes, G. (2012). Action effect anticipation: Neurophysiological basis and functional consequences. Neuroscience and Biobehavioral Reviews, 36(2), 943–959. 10.1016/j.neubiorev.2011.11.004

Winkler, I., Debener, S., Muller, K.-R., & Tangermann, M. (2015). On the influence of high-pass filtering on ICA-based artifact reduction in EEG-ERP. 2015 37th Annual International Conference of the IEEE Engineering in Medicine and Biology Society (EMBC), 4101–4105. d 2.

Wolpe, N., Ingram, J. N., Tsvetanov, K. A., Geerligs, L., Kievit, R. A., Henson, R. N., … & Rowe, J. B. (2016). Ageing increases reliance on sensorimotor prediction through structural and functional differences in frontostriatal circuits. Nature communications, 7(1), 13034. oi.org/10.1109/EMBC.2015.7319296

Zar, J. H. (2010). Biostatistical Analysis (5th ed.). Pearson Prentice Hall.

